# Intertumoral lineage diversity and immunosuppressive transcriptional programs in well-differentiated gastroenteropancreatic neuroendocrine tumors

**DOI:** 10.1101/2022.11.08.515538

**Authors:** Samantha E. Hoffman, Todd W. Dowrey, Carlos Villacorta Martin, Kevin Bi, Breanna Titchen, Shreya Johri, Laura DelloStritto, Miraj Patel, Colin Mackichan, Stephanie Inga, Judy Chen, Grace Grimaldi, Sara Napolitano, Isaac Wakiro, Jingyi Wu, Jason Yeung, Asaf Rotem, Erin Shannon, Thomas Clancy, Jiping Wang, Sarah Denning, Lauren Brais, Ying Huang, Katrina Z. Kao, Scott Rodig, Jason L. Hornick, Sebastien Vigneau, Jihye Park, Matthew H. Kulke, Jennifer Chan, Eliezer M. Van Allen, George J. Murphy

## Abstract

Neuroendocrine tumors (NETs) are rare cancers that may arise in the gastrointestinal tract and pancreas. The fundamental mechanisms driving gastroenteropancreatic (GEP) NET growth remain incompletely elucidated; however, the heterogeneous clinical behavior of GEP-NETs suggests that both cellular lineage dynamics and tumor microenvironment influence tumor pathophysiology. Here, we investigated the single-cell transcriptomes of tumor and immune cells from patients with gastroenteropancreatic NETs. Malignant GEP-NET cells expressed genes and regulons associated with normal, gastrointestinal endocrine cell differentiation and fate determination stages. While tumor and lymphoid compartments sparsely expressed immunosuppressive targets, infiltrating myeloid cells were enriched for alternative immunotherapy pathways including *VSIR*, Tim3/Gal9, and *SIGLEC10*. Finally, analysis of paired primary and metastatic tissue specimens from small intestinal NETs demonstrated transcriptional transformation between the primary tumor and its distant metastasis. Our findings highlight the transcriptomic heterogeneity that distinguishes the cellular landscapes of GEP-NET anatomic subtypes and reveal potential avenues for future precision medicine therapeutics.

## INTRODUCTION

Neuroendocrine tumors (NETs) are rare cancers of the diffuse neuroendocrine system, and while they represent less than 1% of all newly diagnosed malignancies in the United States per year, the population prevalence is estimated to exceed 100,000 individuals (*1*–*3*). The diagnosed incidence of GEP-NETs has risen dramatically over the past four decades, a persistent trend ascribed partially to improved tumor classification and diagnostic technologies (*1, 2, 5*). Approximately two-thirds of all NETs are gastroenteropancreatic (GEP) in origin, predominantly in the pancreas (12-30% of GEP-NETs), stomach, and small intestine (31-50% of GEP-NETs) of older adults (*3, 4*). While patients with localized disease can be cured with surgical resection, these tumors are often identified at a late clinical stage. Despite rapid advancements in NET research and clinical investigation, treatment options for such patients remains limited (*4*).

A limiting factor in identifying novel therapies for neuroendocrine tumors is that the fundamental biology of GEP-NETs remains incompletely understood. Previous investigations have been challenged by limited tissue availability and scarcity of representative *in vitro* and *in vivo* experimental models. Furthermore, genomic mutations in these tumors are most often associated with hereditary endocrine cancer syndromes, including multiple endocrine neoplasia type 1 (*MEN1*), von Hippel-Lindau disease (*VHL*), or tuberous sclerosis (*TSC1/2*) (*6*).

Chromosomal loss, telomere alterations, and epigenetic modifications have emerged as potential regulators of GEP-NET development but are not sufficiently targeted by currently available treatments (*7, 8*). The inter- and intratumoral heterogeneity of GEP-NETs at different anatomical sites (e.g. pancreas, small intestine, stomach) poses an additional, significant challenge to understanding the oncogenic mechanisms driving tumor formation in these disparate environments.

High-resolution delineation of the tumor and immune niches of GEP-NETs has the potential to elucidate novel, actionable targets for future precision medicine endeavors, expanding the limited menu of therapeutic options available to patients with non-resectable tumors. Previous investigations to this end have employed immunohistochemical or bulk RNA-sequencing approaches, but these methodologies are insufficient to identify rare or unknown cell states that significantly contribute to tumor biology and progression. Single-cell RNA-sequencing (scRNA-seq) of other gastrointestinal solid tumors has enabled high-resolution characterization of the cell subpopulations within the cancer microenvironment, thereby overcoming the challenges presented by older technologies (*9*–*12*). We therefore hypothesized that dissecting the transcriptomic landscape of GEP-NETs at single-cell resolution would elucidate shared and distinct cellular phenotypes and genetic programming among these heterogeneous tumors.

## RESULTS

### Characterizing the cellular landscape of gastroenteropancreatic neuroendocrine tumors

We collected fresh surgical resections from 8 patients with well-differentiated NETs, comprising 4 pancreatic NETs (pNETs), 3 small intestinal NETs (siNETs), and one perigastric NET (gNET). Resection tissue was dissociated and divided in two for parallel processing; one half of each cell suspension sorted for viability using flow-cytometry for single-cell RNA-sequencing (scRNAseq) using the 10x Genomics platform, while the other was flow-sorted for both CD45-positivity and viability before undergoing 10x processing **(Figure 1A**, Methods). Of the 8 samples in our cohort, 5 had received no previous therapy, while the remaining 3 tumors had been treated with the somatostatin analogue lanreotide. The gNET had been treated with both lanreotide and chemotherapy with capecitabine and temozolomide (**Figure 1B**). Our cohort also included one primary-metastasis pair from the same patient (sinet2 and sinet3, **Figure 1B**).

**Figure 1.**
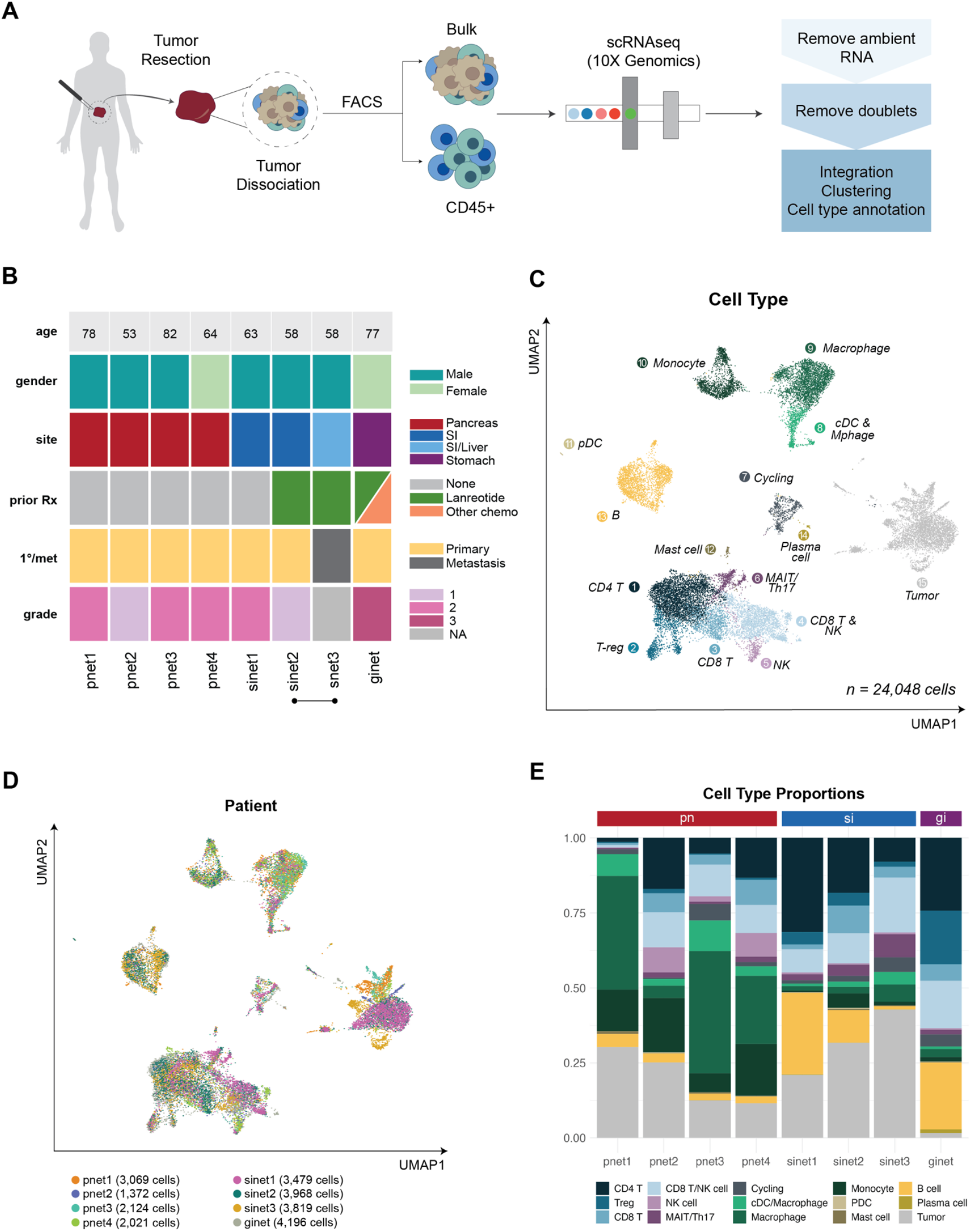
Characterizing the cellular landscape of pancreatic, small intestinal, and perigastric neuroendocrine tumors. **A)** NET single-cell RNA-sequencing workflow. Resected tumor fragments were dissociated into single cell suspensions, which were then divided for parallel sequencing of 1) non-sorted single cells and 2) FACS-sorted, CD45+ immune cells. All samples were processed using the 10X Genomics platform. **B)** Patient demographics and treatment summary at time of tumor resection. Site: SI: small intestine, SI/Liver: small intestinal primary tumor with liver metastasis. A bar underneath samples indicates that they originate from the same patient. **C)** UMAP of malignant and non-malignant cells from all tumors, colored by general cell type. **D)** UMAP of malignant and non-malignant cells from all tumors, colored by patient of origin (left) and inferred presence or absence of malignant copy number variation (CNV, right). **E)** Cellular composition of each tumor, colored by general cell type.

After pre-processing and quality control, our single-cell atlas contained high-quality transcriptomes from 24,048 cells spanning both tumor and immune compartments (**Figure 1C**; Methods). Across-dataset integration of all cells (CD45+-sorted and non-CD45+-sorted) was performed using the *Harmony* algorithm to mitigate batch effects stemming from technical or biological variances between individual patients, tissue collection, and sample processing (**Figure 1C, Figure 1D**; Methods). Using differential gene expression analysis, we identified cluster-specific marker genes that enabled us to subcategorize the lymphoid and myeloid immune compartments (**Figure 1C, Figure S1A**). Tumor cells were identified by high levels of inferred copy number variance relative to non-malignant cell types (**Figure S1C**; Methods).

We then compared the microenvironmental composition of GEP-NETs from distinct sites-of-origin (**Figure 1E**). In general, myeloid cells including macrophages, monocytes, and dendritic cells represented the largest immune subpopulation within pNETs, while siNETs and the gNET contained a relatively higher proportion of T cells and natural killer (NK) cells (**Figure 1E**). Cell-type proportions varied significantly within GEP-NETs from the same site-of-origin, reflecting the intrinsic heterogeneity of these tumors.

### Differential expression analysis and gene regulatory network inference reveal multi-lineage profiles of pNETs

We next interrogated the transcriptional landscapes of GEP-NET cells from different anatomical sites-of-origin (**Figure 2A**). All tumors expressed known neuroendocrine tumor markers such as chromogranin genes *CHGA* and *CHGB* (**Figure 2B**). While most tumors displayed moderate expression of somatostatin receptor 2 (*SSTR2*), no GEP-NET expressed *SSTR5* at significant levels (**Figure 2B**). We also interrogated commonly mutated genes in sporadic pNETs to assess their expression levels across GEP-NETs. *ATRX*, a somatic mutation identified in pNETs, was heterogeneously expressed across GEP-NET subtypes; DAXX, conversely, was either minimally or not expressed by any tumor (**Figure 2B**). Mutations of *ATRX* and *DAXX* in pNETs are often inactivating or missense in nature, which could explain the paucity of expression we identified here (*13*).

**Figure 2.**
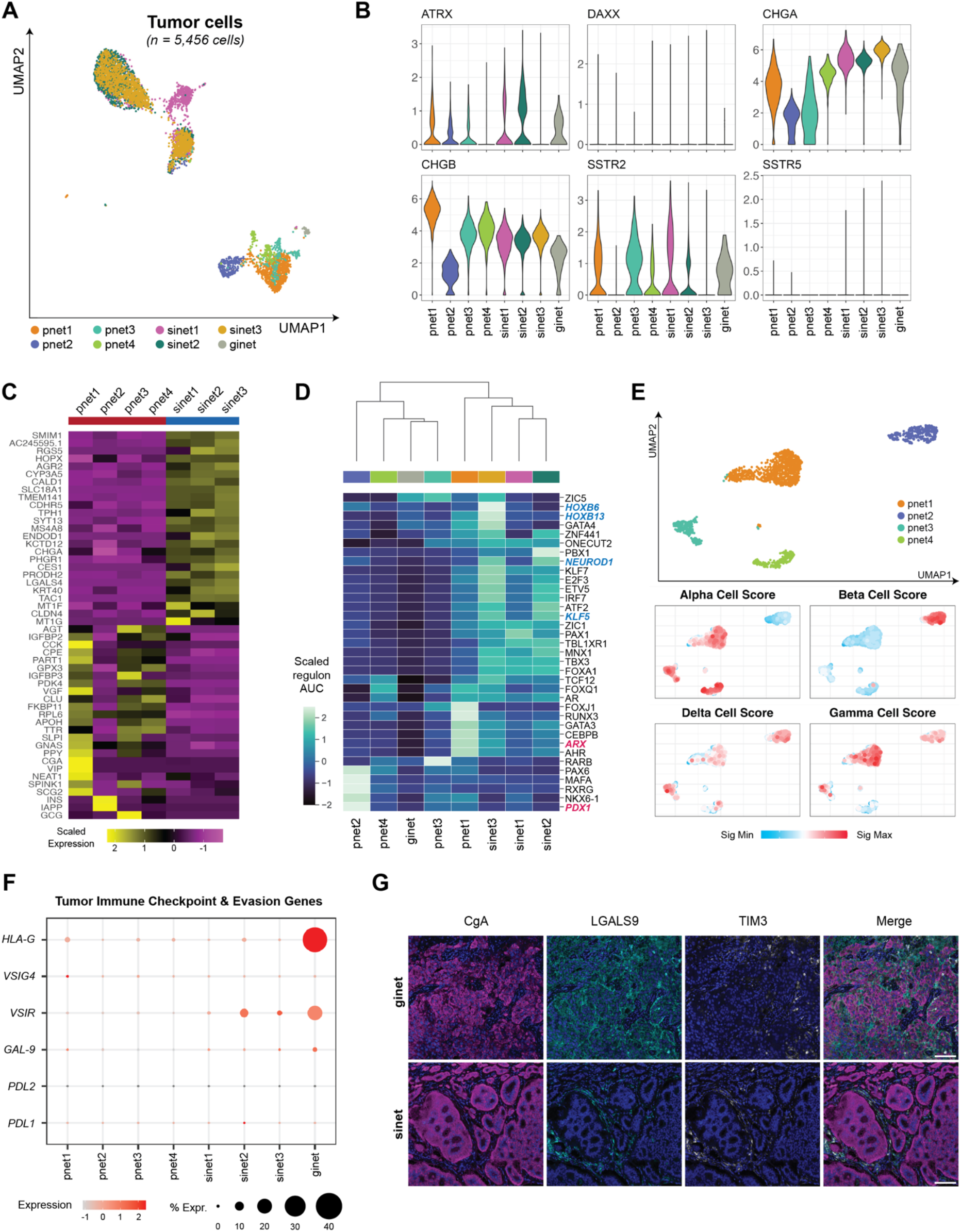
GEP-NET cells do not express common immune checkpoint targets. **A)** UMAP plot of all malignant cells from all GEP-NET samples, colored by patient of origin. **B)** Violin plots of genes commonly expressed in GEP-NETs, grouped and colored by patient. **C)** Heatmap of scaled, normalized average expression of top 50 differentially expressed genes between pancreatic (pn) and small intestinal (si) NETs. **D)** Heatmap of scaled expression of the top five differentially expressed regulons per individual tumor. Blue-labeled genes indicate involvement in enteroendocrine differentiation or regulation of intestinal stem cell fate. Red-labeled genes indicate transcription factors with known involvement in endocrine pancreas development and pNET biology. **E)** UMAP plots of malignant cells from all pNETs, colored by tumor-of-origin (top), followed by heatmaps in UMAP space of VISION signature scores for cell-type specific genes for the major cells in the normal human endocrine pancreas (bottom). **F)** Dotplot of percent expression of common, targetable immune checkpoint and evasion genes, grouped by patient. **G)** Representative immunohistochemical staining of CgA, LGALS9, and TIM3 in one gastric NET and one small intestinal NET sample. LGALS9 and TIM3 expression is largely restricted to the non-tumor compartment. Scale bar represents 100uM.

To compare the transcriptional profiles of pNETs and siNETs, we then performed differential gene expression on all tumor cells from those anatomical sites. Primary and metastatic siNETs exhibited relatively homogeneous transcriptional landscapes compared to pNETs and featured genes involved in monoamine and hormone synthesis (*TAC1, TPH1*, e.g.) and neuroendocrine transport and vesicle release (*SYT13, SLC18A1*, e.g.) (**Figure 2C**), concordant with established profiles of carcinoid tumors. The pNETs in our cohort displayed marked intertumoral heterogeneity, with few differentially expressed genes common to all four tumors (**Figure 2C**). However, all pNETs expressed genes involved in pancreas exocrine (*CCK, SPINK1*, and *SCG2*, e.g.) and endocrine function (*INS, VIP, GCG, PPY*, e.g.) (**Figure 2C**). The neuroendocrine genes expressed by each tumor corresponded closely to its clinical profile; for example, pnet2 demonstrated the highest expression of the insulin-encoding gene *INS*, matching its pathological characterization as a well-differentiated insulinoma (**Figure 2C**).

Next, we sought to identify regulatory elements underlying the heterogeneous molecular profiles of GEP-NETs from different anatomical sites. To this end, we performed single-cell regulatory network inference and clustering (SCENIC) analysis on malignant cells from each tumor. We identified the top regulons for each tumor by AUC value as calculated by the standard pySCENIC workflow (**Figure 2D**, Methods). Hierarchical clustering of NETs by regulon activity produced two major subgroups that did not segregate solely by anatomical location; the first contained most pNETs and the gNET sample, while the other comprised all siNETs and pnet1. Within our pancreatic NET samples, one tumor (pnet2) displayed differential activity of the transcription factors (TFs) *PDX1, PAX6, MAFA, NKX6-1*, and *RXRG* (**Figure 2D**). These TFs are known regulators of beta cell fate in healthy human pancreatic islets, consistent with the clinical diagnosis of insulinoma(*14*–*17*). Another pancreatic NET (pnet1) displayed differential activity of the *ARX* transcription factor, which primarily directs pancreas alpha cell development but also has known roles in establishing pancreatic gamma cell fate in healthy islets of Langerhans (**Figure 2D**)(*18, 19*). These data are consistent with a previous investigation of enhancer profiles of non-functional pNETs, which found that those tumors could be stratified into two major groups by *PDX1* and *ARX* activity(*19*). However, two of our pNETs did not express either *PDX1* or *ARX*. Pnet4 displayed differential activity of *TCF12*, which has been implicated in pancreatic and gastrointestinal cancer proliferation and invasion, while pnet3’s top regulon, *RARB*, was previously found to direct differentiation of embryonic stem cells to pancreatic endocrine cells *in vitro* (*20, 21*). Few common transcription factors were identified across NET samples, revealing the heterogeneity within and between GEP-NETs from different anatomical sites.

Within our siNET samples, we identified differential activity of TFs associated with regulation of enteroendocrine cell lineages (*NEUROD1, FOXA1*) and with the maintenance of intestinal stem cell fate (*KLF5*), concordant with a leading hypothesis in the field of NET biology that siNETs develop from enterochromaffin cells in the gut (**Figure 2D**)(*22*–*24*). The lone siNET metastasis sample (sinet3) displayed highly differential activity of homeobox (*HOX*) transcription factors including *HOXB6* and *HOXB13* (**Figure 2D**). Previous analysis of primary human fetal small intestinal tissue revealed that *HOXB* gene expression is strongest early in duodenal development but quickly diminishes over time (*25*). Neither primary siNET in our cohort displayed significant regulatory activity for any *HOXB* gene, suggesting that the siNET metastasis may have undergone dedifferentiation to resemble an earlier stage of small intestinal fate determination (**Figure 2D**).

To determine the transcriptional similarity of our four pNET tumors to healthy, mature pancreatic islet cells, we scored malignant cells from each tumor with curated gene signatures specific to alpha, beta, gamma, and delta cells in mature human pancreatic islets (**Figure 2E**)(*26*). Tumor cells from pnet1 displayed robust expression of alpha cell and gamma cell gene signatures, with lower expression of the delta cell signature (**Figure 2E**); these findings are consistent with the robust expression of gamma and delta cell hormones *PPY* and *VIP* and elevated *ARX* TF activity identified in our previous analyses. Similarly, as predicted by the results of our differential gene expression and gene regulatory network analysis, pnet2 was the only tumor to demonstrate significant expression of the beta cell signature (**Figure 2E**). Pnet3 and pnet4 scored most highly for the alpha cell signature with lower expression of delta and gamma cell scores (**Figure 2E**). Together with the results of our regulatory network analysis, these data demonstrate tumor cell resemblance to earlier stages of gastrointestinal development as well as the potential for lineage plasticity in these rare tumors.

### GEP-NET cells do not express classical immune checkpoint blockade targets

The success and ongoing studies of immunotherapy in other solid tumor types have motivated a strong interest in applying this therapeutic approach to the treatment of GEP-NETs. We therefore next sought to quantify the expression of classical, tumor cell-anchored immunotherapy targets across the GEP-NETs (**Figure 2F**). Programmed death-ligand 2 (PDL2), which binds programmed cell death protein 1 (PD1) on the surface of T cells to suppress anti-tumoral immunity, was not expressed in any GEP-NETs in our cohort. PDL1, which also binds to PD1 to negatively regulate T cell immunity, was expressed at low levels in fewer than half of tumors. *LGALS9* (also known as Gal-9), *VSIG*, and *VSIR*, other regulators of T cell anergy and exhaustion, were also sparsely expressed but were identified in a greater proportion of tumors than *PDL1*. Multiplex immunofluorescence staining of LGALS9 and its binding partner TIM3 in formalin-fixed, paraffin-embedded tissue from samples in our cohort confirmed low expression of this immunosuppressive axis within the tumor compartment of GEP-NETs (**Figure 2G**). Together, our observations can inform future clinical investigation into the use of anti-PD1 immunotherapies in patients with well-differentiated GEP-NETs.

Given the demonstrated efficacy of the mammalian target of rapamycin (mTOR) inhibitor everolimus in the treatment of patients with advanced GEP-NETs, we also sought to investigate the expression of mTOR pathway genes in our cohort (*27*). We scored all malignant cells with the “PI3K/AKT/mTOR Signaling” gene signature from the HALLMARK collection in MSigDb (**Figure S2A, Figure S2B**) and performed pairwise score comparisons to establish the significance of expression differences (Wilcoxon two-sided test, p < 0.05). The siNETs and giNET in our cohort scored significantly higher for the “PI3K/AKT/mTOR Signaling” signature compared to the pNETs (**Figure S2A**). Interestingly, the metastatic tumor sinet3 had a lower signature score than its primary counterpart (sinet2). Functional follow-up will be required to determine the correlation between gene signature score and everolimus response across GEP-NET subtypes, but our findings demonstrate significant heterogeneity in mTOR signaling across GEP-NETs from different sites-of-origin.

### Profiling lymphoid cell heterogeneity in GEP-NETs

To better understand the immune microenvironment of GEP-NETs, we next analyzed the lymphoid compartment of our dataset. We selected all T and NK cells from the full scRNAseq atlas using canonical marker genes (**Figure S1A**, Methods) for *de novo* clustering and differential gene expression analysis (**Figure S3A, Figure S3B**). We identified diverse T and NK cell populations including NK cells (FGFBP2+ and FGFBP2-), resting and effector CD4+ T cells, regulatory T cells (T-reg), CD8+ T cells, T helper 17 cells (Th17), and mucosal-associated invariant T cells (MAITs) (**Figure 3A, Figure 3B**). These T and NK cell types were identified in differing proportions across GEP-NET site-of-origin, with the gNET containing the highest proportion of T-regs across samples (**Figure 3A**). We also classified B cells from our original dataset, the majority of which originated in either the gNET or siNETs (**Figure S3C**). Of the six B cell clusters identified in our analysis, two contained *IL4R-*expressing naive B cells, while the other four comprised *CD27*-expressing memory B cells (**Figure S3D, Figure S3E**).

**Figure 3.**
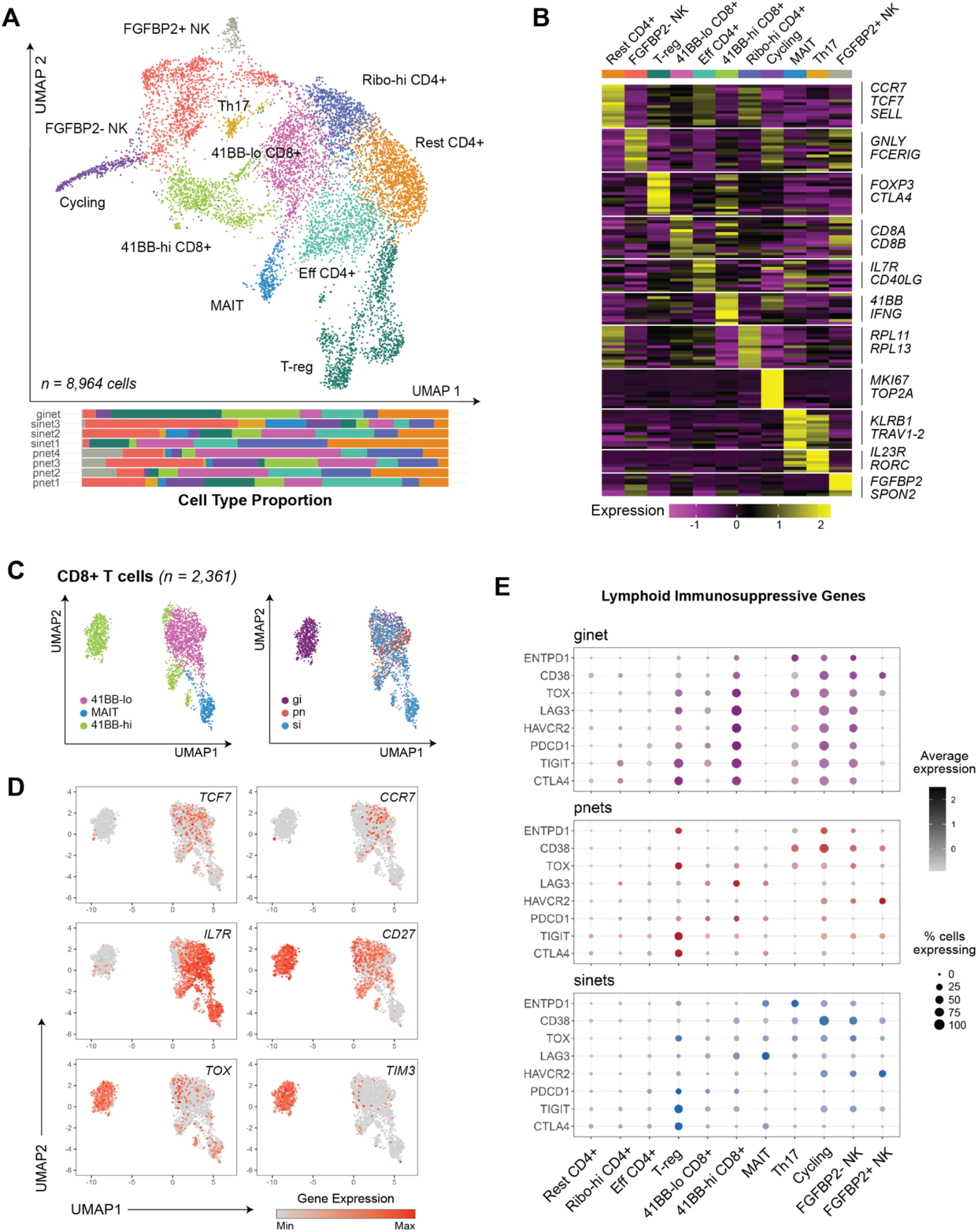
Classifying T cell and NK cell populations within GEP-NETs. **A)** UMAP (top) of T cells and NK cells from all tumors, colored and labelled by lymphoid subtype. Barplot (bottom) displays T/NK cell type proportion by patient. **B)** Heatmap of scaled, normalized expression of top 15 differentially expressed genes per T/NK cell subtype. Specific subtype-determining genes are annotated at right. **C)** UMAP of CD8+ T cells from all tumors, colored by CD8+ T cell subtype (left) and NET origin site (right). **D)** Expression heatmaps in UMAP space of naive (*TCF7, CCR7*), memory (*IL7R*), and exhausted (*TOX*) CD8 T cell markers as well as coinhibitory receptor *TIM3* in CD8+ T cells. **E)** Dotplots of percent of T cell subtypes expressing known immunosuppressive genes, grouped and colored by NET site: gNET: perigastric, pnets: pancreatic, sinets: small intestinal.

The higher expression of immune checkpoint genes in 4-1BB-hi CD8+ T cells compared to other CD8+ T cells across all GEP-NETs motivated us to investigate the exhaustion phenotypes present in infiltrating CD8+ T cells. For this analysis, we included both 4-1BB-hi CD8+ T cells, 4-1BB-lo CD8+ T cells, and MAITs, which demonstrated non-trivial expression of *CD8*. We first isolated all CD8-positive cells from our T and NK cell dataset, which yielded two major subclusters (**Figure 3C**). One cluster contained mostly 4-1BB-hi CD8+ T cells from the gNET sample, while the other cluster represented a mixture of all three CD8+ T cell types from all GEP-NET sites-of-origin. To further delineate CD8+ T cell subtypes, we interrogated the transcription expression patterns of genes previously identified in stem-like T cells (*TCF7*), memory T cells (*IL7R, CCR7*), activated T cells (CD27), and exhausted T cells (*TOX, HAVCR2/TIM3)* (**Figure 3D**). Expression levels of *TCF7* and *IL7R* were highest in MAIT and 4-1BB-lo CD8+ T cells, while *TOX* and *TIM3* displayed the exact opposite expression pattern. These findings are consistent with previous single-cell analyses in renal cell carcinoma, which identified high expression of exhaustion markers among 41BB-hi CD8+ T cells (*28*).

To evaluate the potential therapeutic targets in the GEP-NET milieu using current immunotherapies, we next sought to quantify the expression of known, lymphoid-intrinsic immune checkpoint ligands on the T and NK cells in our atlas. *CTLA4* and *TIGIT* were expressed robustly and frequently in T-regs across all GEP-NET subtypes (i.e. pNETs, siNETs, and gNETs) (**Figure 3E**). *LAG3, HAVCR2* (encoding TIM3), and *PDCD1* (encoding PD1) were sparsely expressed in T and NK cells from pNETs and siNETs. The lymphoid compartment of our one gNET sample demonstrated the highest expression of all queried immunosuppressive markers, with significant upregulation in the 41BB-hi CD8+ T cell, T-reg, FGFBP2-NK cell, and cycling populations (**Figure 3E**). The gNET sample had the highest WHO Grade (Grade 3) of all analyzed samples and was the only tumor to receive treatment with both lanreotide and chemotherapy (**Figure 1B**), which may influence the upregulation of immune checkpoint expression in its T and NK cells. Determining the roles of grade and treatment history in this phenomenon will necessitate expanded serial biopsy cohorts for future analysis.

### Myeloid cells in GEP-NETs express targetable immunosuppressive ligands

Given the limited expression of classical immune checkpoint ligands and receptors on lymphoid cells in the GEP-NET microenvironment, we next sought to characterize the myeloid cells within our cohort. We subset all myeloid cells (n = 4,820 cells) from our original dataset using known marker genes and re-clustered them to identify monocytes (CD16+ and CD16-), dendritic cells (CLEC10A-high DCs, CLEC9A-high DCs, and CD207+ DCs), and tumor-associated macrophages (FOLR2-high TAMs and SELENOP-high TAMs) (**Figure 4A, Figure S4A and S4B**). The proportion of myeloid cell types was similar across all tumors, and we did not identify any patient-specific or GEP-NET subtype-specific myeloid populations (**Figure 4A**).

**Figure 4.**
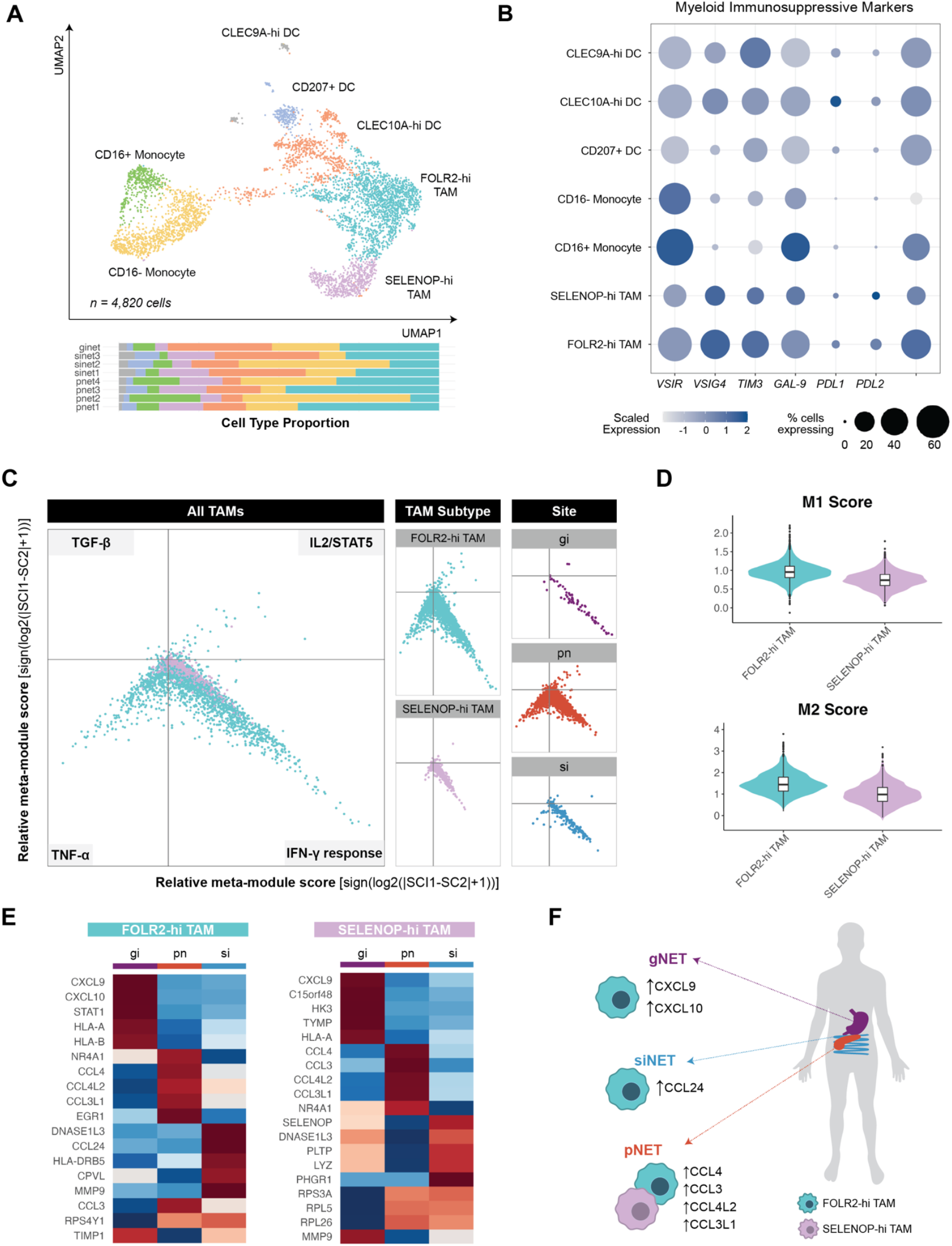
GEP-NET-intrinsic myeloid cells express targetable immunosuppressive ligands. **A)** UMAP of myeloid cells (top) from all tumors, colored and labeled by myeloid cell subtype. Barplot (bottom) displays myeloid cell type proportion by patient. **B)** Dotplot of percentage of each myeloid subtype expressing immunosuppressive ligand genes. **C)** Two-dimensional butterfly plot visualizations of relative meta-module scores of top MSigDB Hallmark Pathways (TNF-α = “TNFA Signaling Via NFKB”, IFN-γ = “Interferon Gamma Response”, IL2/STAT5 = “IL2 STAT5 Signaling”, Complement = “Complement”) in all GEP-NET-associated macrophages (left) and separated by TAM subtype (right). **D)** Violin plot of VISION scores of M1 macrophage- and M2 macrophage-associated gene signatures, grouped by tumor-associated macrophage (TAM) subtype. **E)** Heatmaps of scaled, normalized, and averaged expression of top differentially expressed genes between GEP-NET sites within each TAM subtype (left, FOLR2-hi TAMs and right, SELENOP-hi TAMs). **F)** Graphical summary of differential chemokine expression patterns between TAMs from GEP-NETs at different anatomical sites.

To determine whether myeloid cells in GEP-NETs expressed targetable immune checkpoint ligands, we then quantified the expression of immunosuppressive genes across our nine myeloid clusters. As in the tumor compartment, *PDL1* and *PDL2* were expressed in fewer than 20% of any given myeloid subtype (**Figure 4B**). *VSIG4*, an inhibitor of both cytotoxic T cell and pro-inflammatory macrophage activation, was expressed most strongly in FOLR2-hi TAMs (*29, 30*). Both *TIM3* and its binding partner Galectin-9 were more broadly expressed across both TAM and DC subtypes, suggesting an ability of these cells to suppress the activity of cytotoxic T cells. Finally, the immunosuppressive ligands *VSIR* and *SIGLEC10*, which regulate anti-tumoral macrophage activity, displayed robust expression across the myeloid compartment; *VSIR* specifically was expressed in over 60% CD16+ monocytes and in more than half of SELENOP-hi and FOLR2-hi TAMs (*31*).

To better define the diversity of tumor-infiltrating macrophage phenotypes in GEP-NETs, we isolated the four TAM clusters and performed single-cell pathway enrichment analysis using the HALLMARK genesets from the Molecular Signature Database (Methods). The most significantly enriched pathways included 1) TNF-alpha signaling via NFKB, 2) interferon gamma response, 3) IL2/STAT5 signaling, and 4) genes involved tumor growth-factor beta signaling.

Using the signature scoring protocol outlined in Neftel et. al, we then scored each TAM for expression of each of the four geneset signatures, also referred to as “meta-modules.” We visualized the results of this scoring analysis as a two-dimensional butterfly plot wherein each quadrant represents a specific meta-module (**Figure 4C**, left). FOLR2-hi TAMs scored highly across all four meta-modules, while SELENOP-hi TAMs predominantly expressed TNF-alpha signaling and interferon-gamma response genes. (**Figure 4C**, middle). Grouping TAMs by tumor site-of-origin revealed that while TAMs from pNETs span all four meta-modules, TAMs from siNETs and the gNET predominantly differentially score for the interferon-gamma response signature (**Figure 4C**, right).

Next, we assessed whether GEP-NET TAMs could be defined according to the traditional, albeit debated, M1 vs. M2 macrophage binary classification system (*32, 33*). M1 macrophages are generally considered pro-inflammatory and anti-tumoral, whereas M2 macrophages display immunosuppressive activity. To this end, we scored all TAMs in our cohort for curated gene signatures for M1 or M2 macrophage states (**Figure 4D**). Consistent with similar single-cell analyses in other cancer types, GEP-NET TAMs did not separate neatly into M1 or M2 groupings; for example, FOLR2-hi TAMs scored relatively highly for both the M1 and M2 signatures.

We next sought to determine whether GEP-NET infiltrating TAMs displayed different chemokine profiles depending on the tumor site-of-origin. We performed a differential gene expression analysis within all TAMs in a given subtype grouped by tumor site-of-origin (i.e. pancreatic, small intestinal, or perigastric) and examined the top five differentially expressed genes (**Figure 4E**). All TAMs originating from pNETs displayed increased expression of chemokines *CCL3, CCL4, CCL4L2* and *CCL3L1* (**Figure 4E, 4F)**. FOLR2-hi TAMs from siNETs differentially expressed chemokine *CCL24*, while FOLR2-hi TAMs and SELENOP-hi TAMs in the gNET sample had the highest expression of *CXCL9* and *CXCL10* (**Figure 4E, 4F**). *CCL3, CCL4, CXCL9*, and *CXCL10* are known to recruit activated T cells via binding to the *CXCR3* receptor, suggesting that FOLR2-hi and SELENOP-hi TAM subtypes identified in our cohort may have a role in orchestrating the anti-tumoral immune response in pNETs and gNETs.

*CCL24* has been implicated in activation of the M2 macrophage phenotype; SELENOP-hi TAMs in our siNETs may therefore contribute to a pro-tumoral microenvironment. In sum, our results demonstrate the significant heterogeneity within the myeloid compartments of GEP-NETs and highlight promising therapeutic vulnerabilities common across GEP-NET subtypes.

### Charting the transcriptional heterogeneity and evolution dynamics between a primary siNET and its metastasis

Our cohort contained a matched primary-metastasis pair from a siNET patient, which provided a rare opportunity to investigate the temporal evolution of GEP-NETs. We separated cells from the primary tumor (sinet2) and its liver metastasis (sinet3) for combined analysis (**Figure 5A**). Interestingly, both tumors contained a similar diversity of immune cells despite their different anatomical locations (**Figure 5A**). We then sought to compare the transcriptomic landscapes of the paired samples. Selection of and unsupervised reclustering of only malignant cells from both tumors revealed four tumor clusters (**Figure 5B**). Sinet2, the primary tumor, comprised mostly cells from cluster 0 (C0), with a minority of cells distributed throughout the other three clusters (**Figure 5C**). Conversely, malignant cells from sinet3 fell predominantly in clusters 1 and 2 (C1 and C2); no cells from the metastasis were found in C0 (**Figure 5C**).

**Figure 5.**
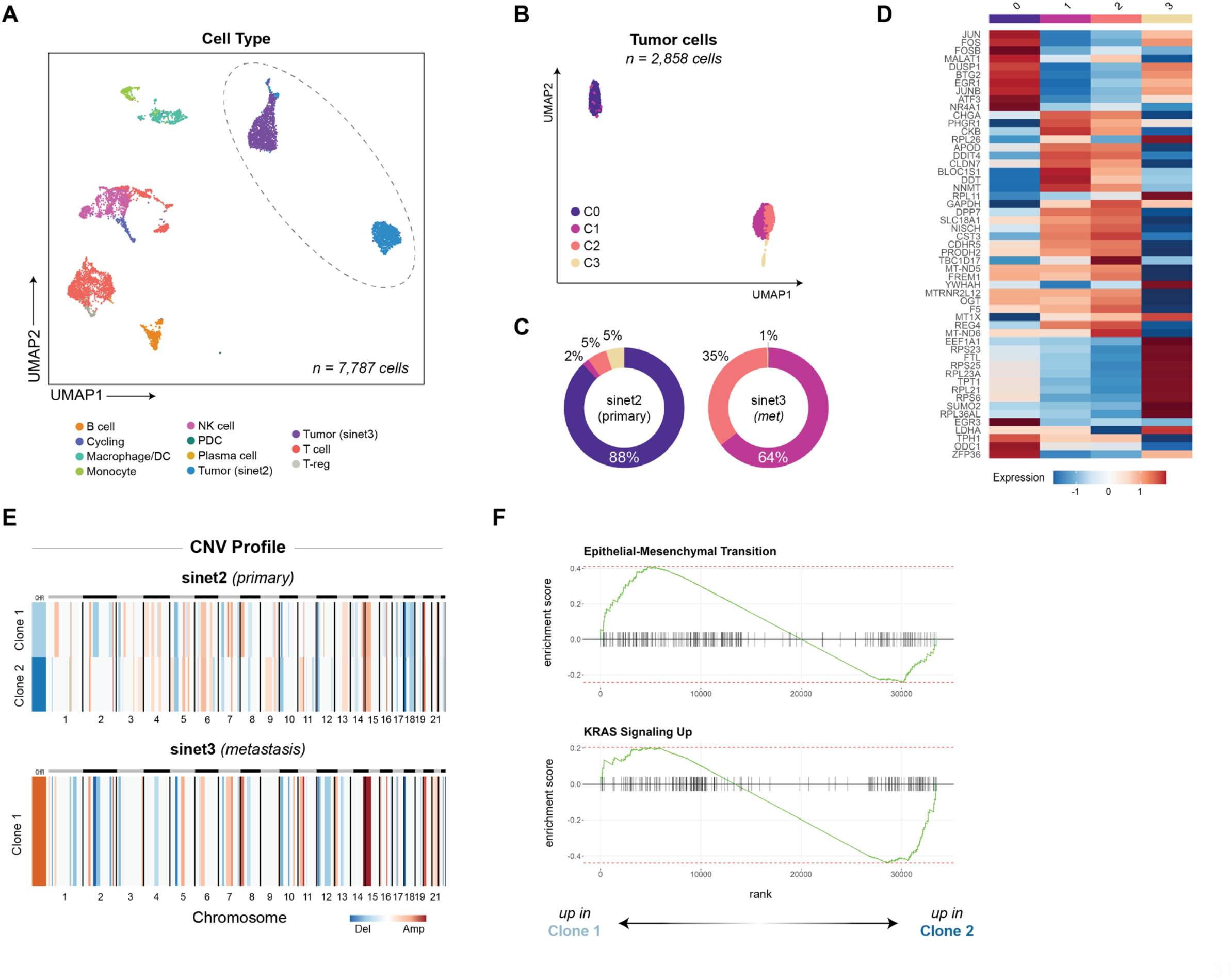
Single-cell RNA-sequencing reveals intratumoral heterogeneity and evolution dynamics in GEP-NETs. **A)** UMAP of malignant and immune cells from a primary small intestinal NET (sinet2) and its paired liver metastasis from the same patient (sinet3). Cells are colored by cancer or immune cell type. Tumor cells are represented by the clusters within the dashed oval. **B)** UMAP of only tumor cells from sinet2 and sinet3, colored by Louvain cluster. **C)** Donut plots representing percentage of cells found in each cluster from Figure 5B for sinet2 and sinet3. Plots are colored by clusters as in Figure 5B. **D)** Heatmap of scaled, normalized, and averaged expression of the top 10 differentially expressed genes between tumor clusters from Figure 5B. **E)** Copy-number alteration profiles of sinet2 and sinet3. Tumor subclones are indicated by color bars on the left-hand side of each plot. **F)** Gene set enrichment plots for the top differential pathways expressed by the two tumor subclones within sinet2.

To interrogate the differences in transcriptional programming between each tumor cluster, we next conducted differential gene expression analysis on the malignant cells from sinet2 and sinet3 (Methods). We identified high expression of immediate early genes such as *JUN, FOS*, and *EGR1* in C0; moderate expression of these genes were also identified in tumor cluster 3 (**Figure 5D**). C1 and C2 were distinguished from the other two clusters by differential expression of neuroendocrine genes such as *CHGA* and *SLC18A1*. Given that C1 and C2 contained mostly cells from sinet3, these data reflect the course of metastatic evolution of sinet2.

To better understand the differential genetic alterations between the primary tumor and its metastasis, we inferred copy number variation profiles for both tumors. The primary tumor (sinet2) contained two clones, with Clone 1 distinguished from Clone 2 by amplifications in chromosomes 1 and 8 as well as deletions in chromosome 2 (**Figure 5E**). The liver metastasis (sinet3) more closely resembled Clone 1, hinting at a possible evolution trajectory between primary tumor and metastatic origin. Next, we sought to interrogate differences in genetic programming between the subclones within sinet2 using gene set enrichment analysis (GSEA, Methods). Clone 1 demonstrated a high enrichment score for the HALLMARK “Epithelial-Mesenchymal Transition” geneset, further implicating this subclone as the likely cellular source of the liver metastasis (**Figure 5F**). Clone 2 was enriched for the HALLMARK “KRAS Signaling Up” geneset (**Figure 5F**). While KRAS mutations have been identified in high-grade pancreatic neuroendocrine carcinomas, further research is required to understand the potential role of KRAS signaling in GEP-NET oncogenesis and progression (*34*). Together, our findings further support the observation of metastatic transcriptional reprogramming in neuroendocrine tumors.

## DISCUSSION

To elucidate the fundamental cellular compositions and genetic programming of GEP-NETs, we interrogated single-cell transcriptomes from eight tumors spanning three anatomical sites-of-origin. Within our findings are reflections of known biology of neuroendocrine tumors. For example, pNETs uniformly demonstrated high differential expression of endocrine pancreatic genes and displayed significantly more intertumoral heterogeneity compared to other NET subtypes. Small intestinal NETs expressed genes associated with monoamine synthesis and neuroendocrine vesicle biology, consistent with the known tendency of these tumors to secrete serotonin. Through scRNA-seq analysis, however, we were also able to further elucidate the transcriptional and gene regulatory network heterogeneity of GEP-NETs. Across our GEP-NET samples, we identified differentially expressed regulons with established roles in neuroendocrine cell differentiation and fate determination that expressed across various timepoints of gastrointestinal development. We further found that the transcriptional profiles of malignant pNET cells resemble more than one cell type in the mature human pancreatic islet including somatostatin-producing delta cells and pancreatic polypeptide (PPY)-producing gamma cells. These data imply the potential for lineage plasticity in GEP-NETs or an endocrine precursor cell provenance for these tumors, two possibilities that should be explored in future biological investigations.

The evolution of neuroendocrine tumors as they metastasize remains poorly understood. While our sample size was limited, we were able to identify two subclones within one of our primary siNETs, one of which bore transcriptomic and copy number alteration similarities to the patient’s liver metastasis. Cells with robust expression of the neuroendocrine genes *CHGA* and *SLC18A1* comprised less than 15% of tumor cells at the primary tumor site but 100% of cells in the metastasis, suggesting that a phenotypic shift occurred over the course of tumor progression and metastasis. Our study is the first to demonstrate the transcriptional dynamics of metastatic neuroendocrine tumors at single-cell resolution; future studies are warranted to investigate the functional and translational significance of the transcriptional programs we identified in our work.

Immunotherapy has revolutionized cancer treatment over the past few decades and has demonstrated benefit across several solid tumor types. Recent clinical trials investigating immunotherapy in neuroendocrine cancers have focused on targeting classical checkpoint genes such as *CTLA4* and the *PD1-PDL1/PDL2* axes in the lymphoid compartment (*35, 36*).

While many of these studies are ongoing, preliminary results suggest that undifferentiated, higher-grade neuroendocrine neoplasms may respond to checkpoint inhibition, but that use of checkpoint inhibitors in well-differentiated NETs, like those in our study, would yield limited therapeutic benefit (*37*–*40*). Consistent with these clinical observations, within all GEP-NET cancer cells in our dataset, we identified little to no expression of currently targetable immune checkpoint ligands, highlighting the limited treatment options in this disease. Similarly, within the lymphoid compartment of GEP-NETs, we further identified relatively low expression of common immunosuppressive ligands and receptors across T and NK cell types. 4-1BB-hi CD8+ T cells and FGFBP2-NK cells from our perigastric NET exhibited the highest expression of immune checkpoint genes; however, this sample was the only higher grade (grade 3) tumor of our cohort and the only one previously exposed to chemotherapy, both factors that could have contributed to this finding. Further translational studies will be required to determine the clinical utility of targeting lymphoid cells with immunotherapy in well-differentiated NETs.

In contrast, within the myeloid compartment, we observed high levels of immunosuppressive gene expression in multiple cell types across GEP-NET sites-of-origin. Our data highlight opportunities for therapeutic strategies in these rare cancers based on targeting alternative pathways in the myeloid microenvironment of GEP-NETs. For instance, high expression of *VSIR* across myeloid cells in GEP-NETs suggests that targeting the VISTA protein could rearm T cells and macrophages and thereby reinvigorate anti-tumoral immunity in these patients. Preclinical studies of the VISTA pathway in multiple solid tumor models (pancreatic cancer, melanoma, colorectal cancer, renal cell carcinoma, and glioma, e.g.) have affirmed its role in tumoral immune evasion (*31, 41, 42*). Early clinical trial data are assessing the safety profiles of anti-VISTA immunotherapies in solid tumors that also target the PD-1/PD-L1 checkpoint; per our findings, well-differentiated NETs may benefit from this therapeutic approach and should be considered for inclusion in future trial cohorts (*43, 44*). Similarly, we identified significant expression of *HAVCR2* (TIM3) and *LGALS9* (Gal-9) in myeloid cells of the GEP-NET microenvironment. Combination anti-Tim3/anti-PD-1 or anti-Gal-9/anti-PD-1 therapies have proven efficacious in enhancing T cell activation in preclinical studies of solid tumors and are currently under investigation in clinical trials (*45*–*47*). Finally, inhibitors of SIGLEC10-CD24 binding between TAMs and cancer cells, respectively, are an emerging focus of preclinical investigation (*48*). Our study therefore provides significant evidence supporting the use of these promising, emerging immunotherapies, which may one day augment the limited effective treatment options available to patients with non-resectable GEP-NETs.

Our study has several limitations While the diversity of our cohort permits us to assess shared and common transcriptional programs across GEP-NET subtypes, our sample size remains relatively small due to the rarity of these tumors. Treatment history at the time of surgical resection also varied between sites-of-origin; all pancreatic NETs were treatment-naive, while most of the small intestinal and gastric NETs had received at least lanreotide therapy.

Differences in cellular phenotypes and microenvironment composition may therefore reflect therapeutic pressures and not necessarily innate biology. Nevertheless, our cohort encompasses multiple, well-differentiated GEP-NET subtypes across diverse treatment histories; our findings therefore serve as a key foundation for future basic and clinical investigations into these rare, heterogeneous tumors.

In summary, our findings uncover previously unappreciated heterogeneity within neuroendocrine tumor subtypes and reveal potential evolution in tumor characteristics as they metastasize. Consistent with other studies, we also found that, in well differentiated neuroendocrine tumors, expression of checkpoint markers targetable with current standard immunotherapies is uncommon. However, we also identified new immunosuppressive markers in myeloid cells that are commonly expressed in neuroendocrine tumors and may form the basis for future targeted therapies. Our findings therefore provide an important step in delineating key underlying molecular features of neuroendocrine tumors and shed light on potential novel therapeutic strategies for patients suffering from this challenging disease.

## METHODS

### Lead Contact

Further information and requests should be directed to the corresponding authors.

### Data and Code Availability

Raw single-cell RNA-seq data will be deposited into dbGaP. Processed single-cell RNA-seq data will be deposited into the Broad Institute Single Cell Portal (https://singlecell.broadinstitute.org/single_cell). We will include annotations for all cells including patient-of-origin, tissue site, and cell-type metadata used for this manuscript’s analyses. Code for analysis is described in “Method Details” below.

### Experimental Model and Subject Details

Human patient samples were collected under Dana-Farber Cancer Institute IRB protocol #02-314.

### Methods Details

#### Sample collection and dissociation for scRNA-seq

Fresh surgical resections were collected at Brigham and Women’s Hospital. Tumor tissue was dissected away from non-tumor tissue using a scalpel, transferred to a 50 mL conical tube with HBSS and transported on ice. On arrival to the laboratory, tumor samples were transferred into a 5mL Eppendorf tube containing 4.5mL cold enzymatic dissociation mix (100µg/mL Collagenase Type 4 (Worthington Biochemical Corporation #LS004186) and 10 µg/mL DNAse I (StemCell Technologies #07900) in RPMI + HEPES (ThermoFisher Scientific #22400089)). The resection tissue was then minced inside the tube with spring scissors into <0.5mm fragments at room temperature. The resulting mixture was incubated for 10 minutes in a 37°C water bath with manual inversions every 1-2 minutes to ensure thorough mixing. Next, the mixture was vigorously and repeatedly pipetted using a P1000 pipette at room temperature for further mechanical dissociation. For all samples except pnet2 and pnet3, the tube containing the mixture was then returned to the 37°C water bath for an additional 10 minutes with manual inversions every 1-2 minutes for thorough mixing.

The cell suspension was then filtered through a 70µm cell strainer into a 15mL conical tube on ice. The strainer was washed with 5mL cold RPMI to maximize cellular yield. The filtered cells then underwent centrifugation at 450xg for 5 minutes at 4°C. The resulting supernatant was carefully transferred to a 15mL conical tube on ice, taking care not to disturb the pellet. To lyse red blood cells, the centrifuged cell pellet was resuspended in 300-500µL of ACK Lysing Buffer (ThermoFisher Scientific #A1049201) and incubated for 1 minute on ice.

Calcium- and magnesium-free PBS (ThermoFisher Scientific #10010-23) was added in a volume twice that of the ACK Lysing Buffer to stop red blood cell lysis. Cells were transferred to a 1.7mL Eppendorf tube on ice and then recentrifuged for 8 seconds at 4°C with centrifugal force ramping up to but not exceeding 11,000xg. After re-centrifugation, cells were resuspended in calcium- and magnesium-free PBS with 0.4% BSA (Ambion #AM2616). If a significant amount of red blood cells remained after recentrifugation, an additional round of red blood cell lysis was performed. After satisfactory red blood cell lysis, the supernatant was carefully transferred to a new 15mL conical tube on ice, and the remaining cell pellet was resuspended in 50µL of RPMI + HEPES.

5µL of the single-cell suspension was mixed with Trypan blue (Sigma #T8154) at a 1:1 volume ratio and loaded into a hemocytometer for cell counting and assessment of cell viability, clumping, and debris. At the end of processing, all samples had viability greater than 51%. Cell suspensions were then centrifuged and resuspended in either RPMI + HEPES or RPMI + HEPES with 0.4% BSA in preparation for transport and downstream flow cytometric processing.

#### Flow cytometry

Following viability assessment, samples were centrifuged at 600xg for 5 minutes and resuspended in FACS buffer (1% BSA (Millipore Sigma, Product No: EM-2930) in calcium/magnesium-free PBS (ThermoFisher; Product No: 14190144)) and stained with 1 µg CD45-APC antibody (Fisher Scientific; Product No: 50-166-056) for 30 minutes at 4°C. The samples were washed with 1 mL FACS buffer and centrifuged at 600xg for 5 minutes before being resuspended in 100 µL FACS buffer with 10 µM Stemolecule™ Y27632 (ROCK inhibitor) (Stemgent; Product No: 04-0012-02). 200 µL FACS buffer with calcein blue AM (Fisher Scientific; Product No: C1429) was added to the sample for live/dead exclusion and the sample was passed through a 35 µm strainer prior to sorting. Two fractions of the samples (live and CD45^+^) were sorted into FACS buffer using a Beckman Coulter MoFlo Astrios cytometer.

#### Single-cell RNA-sequencing

For each sample, the live bulk and CD45^+^ fractions were centrifuged at 600xg for 5 minutes and resuspended in FACS buffer for a final concentration of 600 cells/µL as specified in the 10x Genomics user guide document number CG000185, Rev B for an estimated recovery of 3000 cells. These fractions were loaded into the 10X Genomics Single Cell chip (10X Genomics # PN-1000120) along with gel beads and reverse transcription master mix from the Single Cell 5’ v1.1 Chemistry Kit (10x Genomics # PN-1000165). The chip was run on the 10X Chromium Controller using the standard program to produce cDNA, which was then amplified and used for Gene Expression (GEX) library generation. All single cell GEX libraries were sequenced using an Illumina NextSeq 2000 sequencer.

#### scRNA-seq data preprocessing

Raw sequencing data underwent demultiplexing, barcode processing, read alignment, and UMI counting using the 10X Genomics CellRanger pipeline (version 5.0) with default parameters. Reads were aligned to the pre-built human genome reference included in Cell Ranger (GRCh38). After filtering to exclude low-RNA content droplets, a gene-barcode matrix was generated for each sample containing counts of confidently mapped, non-PCR-duplicate reads.

To remove technical artifacts arising from cell-free RNA profiles from our sequencing data, ambient RNA decontamination was performed on all samples using the CellBender software package (*49*). In brief, the raw counts matrix and expected cell count for each sample was provided as input to the *remove-background* module of CellBender, which employs an unsupervised deep-generative model to distinguish empty droplets from cell-containing ones. Expected cell count was derived from the estimate generated by Cell Ranger. All samples were run using the “ambient” model and default settings of 150 training epochs and false positive rate of 0.01. Transcripts assigned to empty cells were iteratively detected and removed from the raw counts matrix by *remove-background*, yielding a “background-subtracted” cleaned counts matrix as output.

To remove multiplet artifacts caused by droplet encapsulation of more than one cell, we then performed doublet detection and removal on the CellBender-cleaned counts matrix using the Scrublet package (Python, version 0.2.2) (*50*). An expected doublet rate of 0.06 and manually selected doublet score thresholds were used to separate singlets from neotypic doublets in the resulting bimodal score distribution graphs. Presumed doublets were removed to create an ambient RNA-free, singlet-only cleaned counts matrix for downstream analysis. All further quality control, dimensionality reduction, unsupervised clustering, and differential expression analyses were performed using the Seurat R package (version 4.0.6)(*51*).

#### Sample quality control and integration

Cells with fewer than 200 genes or greater than 25% counts representing mitochondrial genes were removed from each sample. Genes detected in fewer than three cells per sample were also removed. The raw counts data was then normalized and scaled using the default log-normalization approach from the R package Seurat (version 4.0.6) (*51*). To mitigate technical and biological variation between samples, the sixteen datasets (one unsorted and one CD45+-sorted per sample) were integrated using the R package Harmony (version 0.1.0) using patient-of-origin as the primary source of variation (*52*). The final integrated dataset included 24,048 cells from the eight-tumor cohort.

#### Cell type clustering, visualization, and identification

Linear dimensional reduction was performed on the integrated and normalized counts matrix by running principal component analysis on the 5000 most highly variable genes in our dataset. The first 30 principal components were used to perform Louvain clustering with a resolution parameter of 0.4. Uniform Manifold Approximation and Projection (UMAP) was then performed using the same principal components to visualize cell clusters in two-dimensional space. To determine cell types within each cluster, differential gene expression analysis was performed by comparing cells from each cluster to all other cells using a two-sided Wilcoxon ranked-sum test with applied Bonferroni correction for multiple comparison testing. Top differential genes within each cluster were used to assign cells into broad tumor and immune categories. For subclassification of immune cell types, lymphoid and myeloid cells were subset from the integrated dataset and clustered and UMAP-projected using 30 principal components. The same differential gene expression analysis was used to identify T cell and myeloid types at higher resolution. CD8+ T cells and tumor-associated macrophages were further re-clustered and re-projected to conduct similar cellular dissection.

#### Identification of cancer cells

To identify malignant cells, we first integrated both non-CD45+-sorted and CD45+-sorted datasets from the same tumor to generate a combined expression matrix. Each sample underwent Louvain clustering and UMAP-projection to identify *PTPRC+* (CD45+) and *PTPRC-*clusters. We then used inferCNV (R, Terra implementation) to estimate the CNV profile of each log-normalized sample using all *PTPRC+* cells as the reference group and all *PTPRC-*cells as the observation group. NET-related marker gene expression and malignant calls from inferCNV were combined to identify “Tumor” cells within our cohort.

#### Differential expression, gene set enrichment analysis, and signature scoring

To minimize the effect of patient-specific expression patterns on our differential gene expression analysis of GEP-NET cells across anatomical subtypes (**Figure 2C**), we averaged the expression of genes from all tumors from the same primary site-of-origin (i.e. pancreas or small intestine) and compared the two transcriptional profiles. To compare the transcriptomic profiles of cells from a paired primary (sinet2) and metastatic (sinet3) tumor set, differential gene expression analysis was performed using a two-sided Wilcoxon rank-sum test with Bonferroni correction applied. The log_2_(fold-change) values for each gene were used as ranks for preranked gene-set enrichment analysis (GSEA), which was performed using the fgsea R package (version 1.20.0) (*53*). GSEA was performed using all HALLMARK, KEGG, and GO Biological Process gene sets from version 6.2 of the Molecular Signatures Database (MSigDb) as input (*54*). Significance level was set at p < 0.05 and FDR < 0.25.

Single-cell signature scoring was performed using the VISION R package (version 3.0.0) (*55*). Progenitor exhaustion and terminal exhaustion signatures derived from Bi et al, Cell 2021 were used to score CD8+ T cells from all tumors (Figure 3F). Known macrophage, monocyte, and dendritic cell markers were used to generate custom gene signatures, which were then used for VISION scoring of the NET myeloid compartment (Figure 3B).

#### SCENIC analysis and regulon inference

To identify differentially active gene regulatory networks (GRNs) in tumor cells across GEP-NETs, we performed single-cell regulatory network inference and clustering (SCENIC) analysis using the recommended pipeline provided with the pySCENIC implementation ((http://htmlpreview.github.io/?https://github.com/aertslab/SCENICprotocol/blob/master/notebooks/PBMC10k_downstream-analysis.html) (*56, 57*). We first performed GRN inference using the multiprocessing implementation of Arboreto to run GRNBoost2 on the count matrix containing averaged expression of tumor cells from each sample. We then ran the SCENIC implementation of cisTarget to predict regulons from the transcription factor adjancencies matrix produced by GRNBoost2. Finally, we used AUCell to generate regulon specificity scores for each tumor. As recommended in the pySCENIC documentation, we selected the top 5 regulons in each tumor ranked by AUC, then plotted the scaled scores for these regulons by tumor.

#### Meta-module scoring

To generate meta-module scores for tumor-associated macrophages in Fig. 4, we first used Seurat’s FindAllMarkers function to identify TAM cluster-specific marker genes. We then used the average log_2_(fold-change) of genes in our differential expression analysis as a ranking metric for GSEA analysis on all TAMs using the HALLMARK genesets from the Molecular Signatures Database (MSigDB). Four of the most significantly enriched pathways from this analysis were selected as biologically relevant meta-modules. To score each TAM for expression of these four meta-modules, we implemented the method developed in Neftel et al. (*58*) using the scrabble R package. Briefly, a signature score (SC) is calculated for each cell by averaging the relative expression of the signature genes in that cell, then subtracting the average relative expression of a control geneset in the same cell. The control geneset is defined as previously (*58*). All cells were scored for each of the four meta-modules. The position of each cell in the butterfly plot (Figure 4C) was determined by calculating [sign(SC1-SC2)*log_2_(|SC1-SC2|+1)] for both x- and y-coordinates.

#### Determination of CNV clonal structure

To determine the clonal variation between the primary (sinet2) and metastatic tumor cells (sinet3) from the same patient with a small intestinal NET, cells from both samples were processed using the *pipelineCNA* function from the R package SCEVAN (*59*). The “subclones” parameter was set to “TRUE” to enable analysis of clonal structure within each tumor.

Heatmaps displaying inferred copy number loss/gain were generated for each sample and then compared to identify changes in clonal substructure between the primary tumor and its metastasis.

#### Multiplex immunofluorescence staining and imaging

To confirm the expression of immunosuppressive markers identified by our computational analysis, formalin-fixed, paraffin-embedded (FFPE) slides from each tumor in the cohort were stained with both DAPI and two, six-marker multiplex immunofluorescence panels. Two seven-plex Fluorescence Immunohistochemistry assays were performed on consecutive 4-µm FFPE sections using the Leica Bond Rx autostainer. The immune panel consisted of six antibodies: CD45 (2B11 + PD7/26, Mouse monoclonal, Dako) at 1:200, CD3 (Rabbit polyclonal, Dako) at 1:100, CD4 (EP204, Rabbit monoclonal, Sigma-Aldrich) at 1:250, CD8 (C8/144B, Mouse monoclonal, Dako) at 1:1,000, CD68 (PG-M1, Mouse monoclonal, Dako) at 1:100, FOXP3 (D2W8E, Rabbit monoclonal, Cell Signaling) at 1:250; The NET panel included six antibodies: Chromogranin A (DAK-A3, Mouse monoclonal, Dako) at 1:200, Synaptophysin (DAK-SYNAP, Mouse monoclonal, Dako) at 1:200, LGALS9 (OTI19H8, Mouse monoclonal, Origene) at 1:200, TIM3 (EPR22241, Rabbit monoclonal, Abcam) at 1:1,000, PD1 (EH33, Mouse monoclonal, Cell Signaling) at 1:200, PD-L1 (E1L3N, Rabbit monoclonal, Cell Signaling) at 1:200. Briefly, the staining consists of sequential tyramine signal amplified immunofluorescence labels for each target, along with a DAPI counterstain. Each labeling cycle consists of application of a primary antibody, a secondary antibody conjugated to horse radish peroxidase (HRP), and an opal fluorophore (Opal 520, Opal 540, Opal 570, Opal 620, Opal 650 and Opal 690, Akoya Biosciences), respectively.

The stained slides were scanned on a Perkin Elmer Vectra 3 imaging system (Akoya Biosciences) and analyzed using Halo Image Analysis platform (Indica Labs). Each single stained control slide is imaged with the established exposure time for creating the spectral library. We ran an algorithm learning tool utilizing the Halo image software training for the gland and stroma regions, and subsequently completed cell segmentation. The thresholds for the antibodies were set respectively, based on the staining intensity, by cross reviewing more than 20 images. Cells with the intensity above the setting threshold were defined as positive.

## ACKNOWLEDGEMENTS

We thank the patients who participated in this study. We are grateful to K. Pfaff and the Center for Immuno-Oncology at Dana-Farber Cancer Institute for assistance with the interpretation of immunofluorescence in patient samples.

## Funding

This work was supported by:

National Institutes of Health grant T32 GM141745 (S.E.H)

National Institutes of Health grant R37 CA222574 (J.P., E.M.V.A)

National Institute of Health grant R01 CA227388 (J.P., E.M.V.A)

Bristol Myers Squibb (S.R.)

KITE/Gilead (S.R.)

## Author information

**Colin Mackichan**

Present address: Viome, Bothell, WA, USA

**Judy Chen**

Present address: Yale School of Public Health, New Haven, CT, USA

**Grace Grimaldi**

Present address: Memorial Sloan Kettering Cancer Center, New York, NY, USA

**Isaac Wakiro**

Present address: Memorial Sloan Kettering Cancer Center, New York, NY, USA

**Jingyi Wu**

Present address: University of Rochester, Rochester, NY, USA

**Jason Yeung**

Present address: Boston University, Boston, MA, USA

**Asaf Rotem**

Present address: AstraZeneca, Waltham, MA, USA

**Ying Huang**

Present address: Novartis Institutes for BioMedical Research, Cambridge, MA, USA

**Sebastien Vigneau**

Present address: Ginkgo Bioworks, Boston, MA USA

## AUTHOR CONTRIBUTIONS

S.E.H: conceptualization, formal analysis, writing – original draft, visualization, methodology T.W.D.: methodology, investigation, writing – review & editing

C.V.M.: data curation, formal analysis

K.B.: writing – review & editing

B.T.: writing – review & editing

S.J.: writing – review & editing

L.D.: data curation, project administration

M.P.: investigation

C.M.: investigation

S.I.: investigation

J.C.: investigation

G.G.: investigation

S.N.: investigation

I.W.: investigation

J.Wu.: investigation

J.Y.: investigation

A.R.: investigation

E.S.: project administration

T.C.: resources

J.W.: resources

S.D.: investigation

L.B.: project administration

Y.H.: data curation, formal analysis

K.Z.K.: data curation, investigation

S.R.: supervision

J.H.: resources

S.V.: investigation, supervision

J.P.: writing – review & editing

M.H.K.: conceptualization, funding acquisition, supervision, writing – review & editing

J.C.: conceptualization, funding acquisition, supervision, writing – review & editing

E.M.V.A: conceptualization, funding acquisition, supervision, writing – original draft, writing – review & editing

G.J.M.: conceptualization, funding acquisition, supervision, writing – review & editing

## DECLARATION OF INTERESTS

AR is an equity holder in Celsius Therapeutics and NucleAI. J.C. reports consulting relationships with Advanced Accelerator Applications and TerSera and equity in Merck. E.M.V.A reports advisory and consulting relationships with Tango Therapeutics, Genome Medical, Genomic Life, Enara Bio, Janssen, Manifold Bio, and Monte Rosa; research support from Novartis and BMS; equity in Tango Therapeutics, Genome Medical, Genomic Life, Syapse, Enara Bio, Manifold Bio, Microsoft, and Monte Rosa; institutional patents filed on chromatin mutations and immunotherapy response, and methods for clinical interpretation; intermittent legal consulting on patents for Foaley & Hoag; and serves on the Editorial Boards of *JCO Precision Oncology* and *Science Advances*. All other authors declare they have no competing interests.

## SUPPLEMENTARY FIGURE LEGENDS

**Figure S1.**
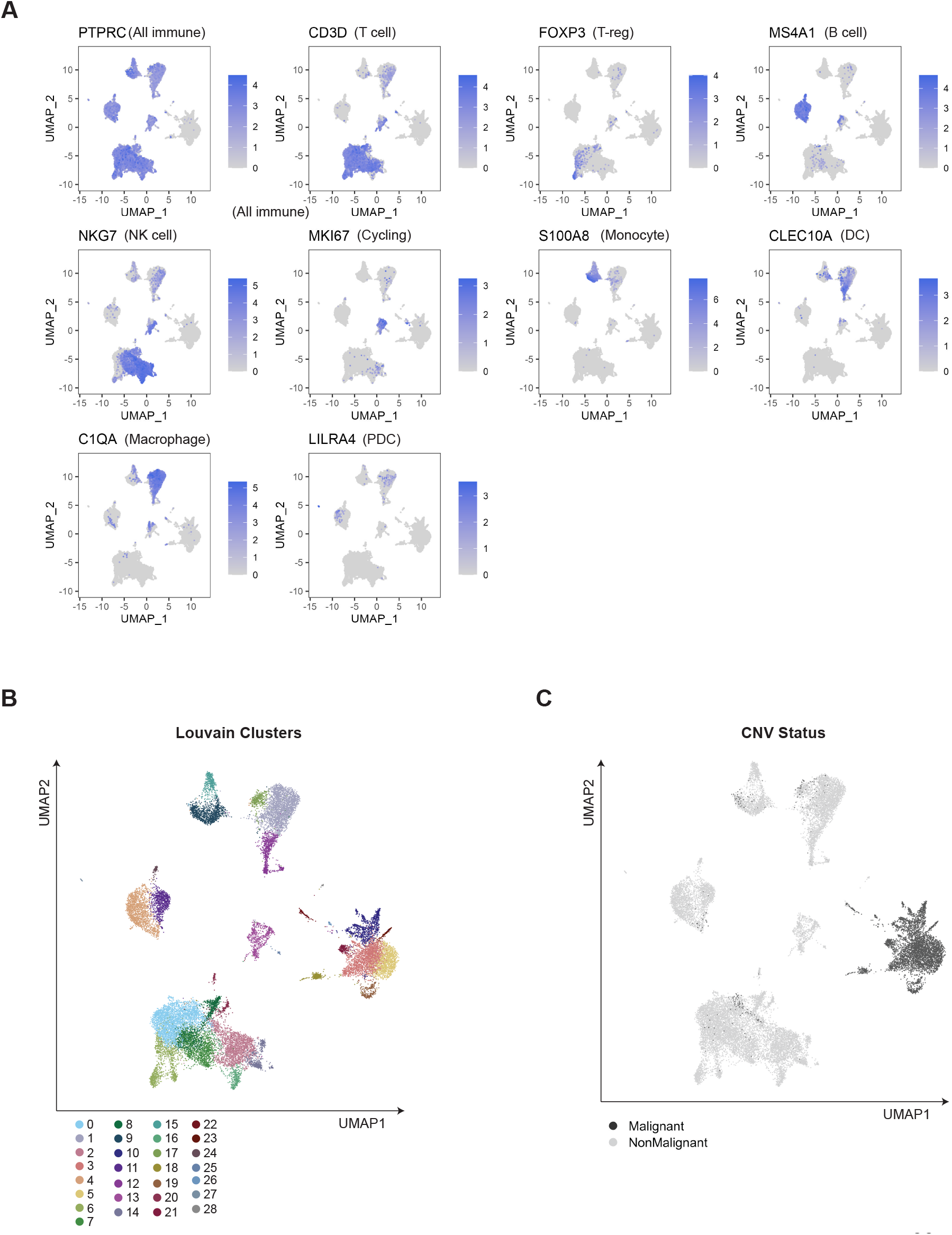
Identification of general tumor and immune cell types and sub-lineages. Related to Figure 1. A) Expression UMAP plots of malignant and non-malignant cells from all tumors representing expression of one marker gene per general immune cell type. B) UMAP of all malignant and non-malignant cells from all GEP-NET samples, colored by initial Louvain clustering. C) UMAP of all malignant and non-malignant cells from all GEP-NET samples, colored by presence or absence of copy-number variation as inferred by *inferCNV* analysis.

**Figure S2.**
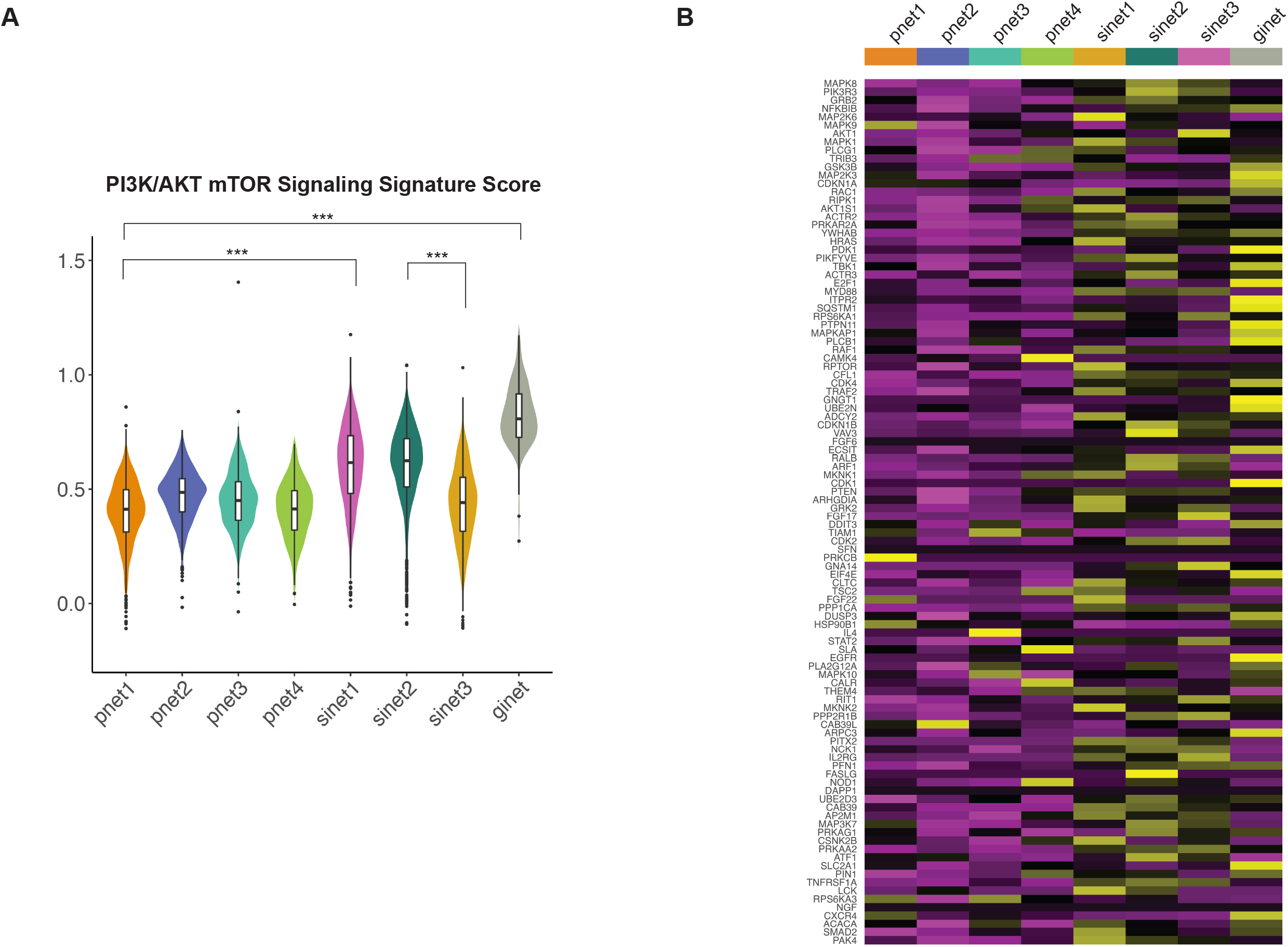
Related to Figure 2. A) Violin plots of HALLMARK PI3K/AKT/mTOR signature score distributions in malignant GEP-NET cells, separated by tumor-of-origin. Significance of signature score enrichment was calculated using a two-sided Wilcoxon rank-sum test (*** = p < 0.001). B) Heatmap of scaled, normalized, and averaged expression 105 genes in the HALLMARK “PI3K/AKT/mTOR Signaling” signature on a per-tumor basis.

**Figure S3.**
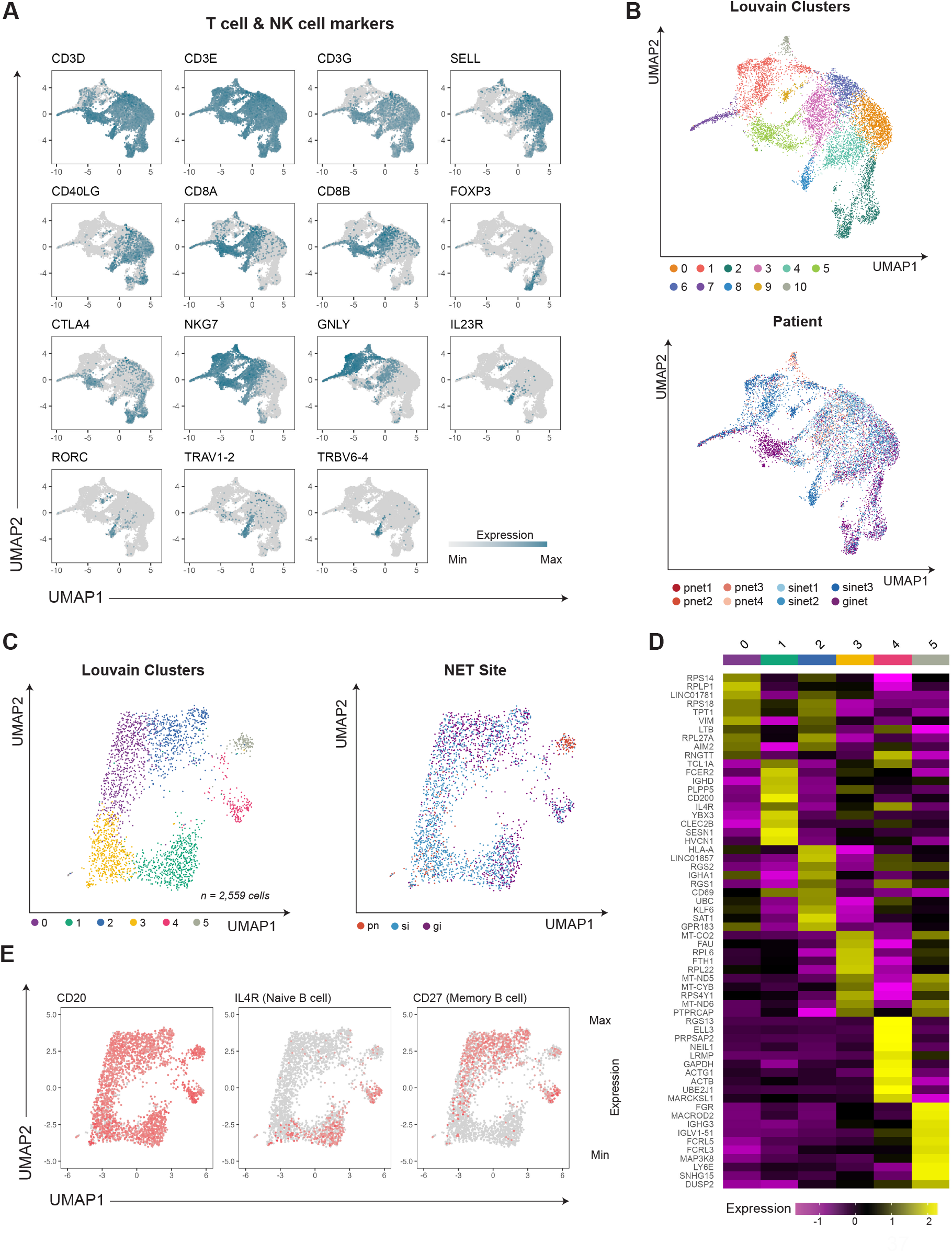
Related to Figure 3. A) Expression UMAP plots of T and NK cells from all tumors, representing expression of marker genes used to identify lymphoid subtypes. B) UMAP of all T and NK cells from all GEP-NET samples, colored by initial Louvain clustering (top) and patient (bottom). C) UMAP of all B cells and plasma cells from all GEP-NET samples, colored by initial Louvain clustering (left) and GEP-NET site-of-origin (right). D) Heatmap of scaled and normalized average expression of top 10 differentially expressed genes between B cell and plasma cell clusters. E) Expression UMAP plots of B cells from all tumors, representing expression of pan-B cell marker CD20 (left), naïve B cell marker IL4R (middle), and memory B cell marker CD27 (right).

**Figure S4.**
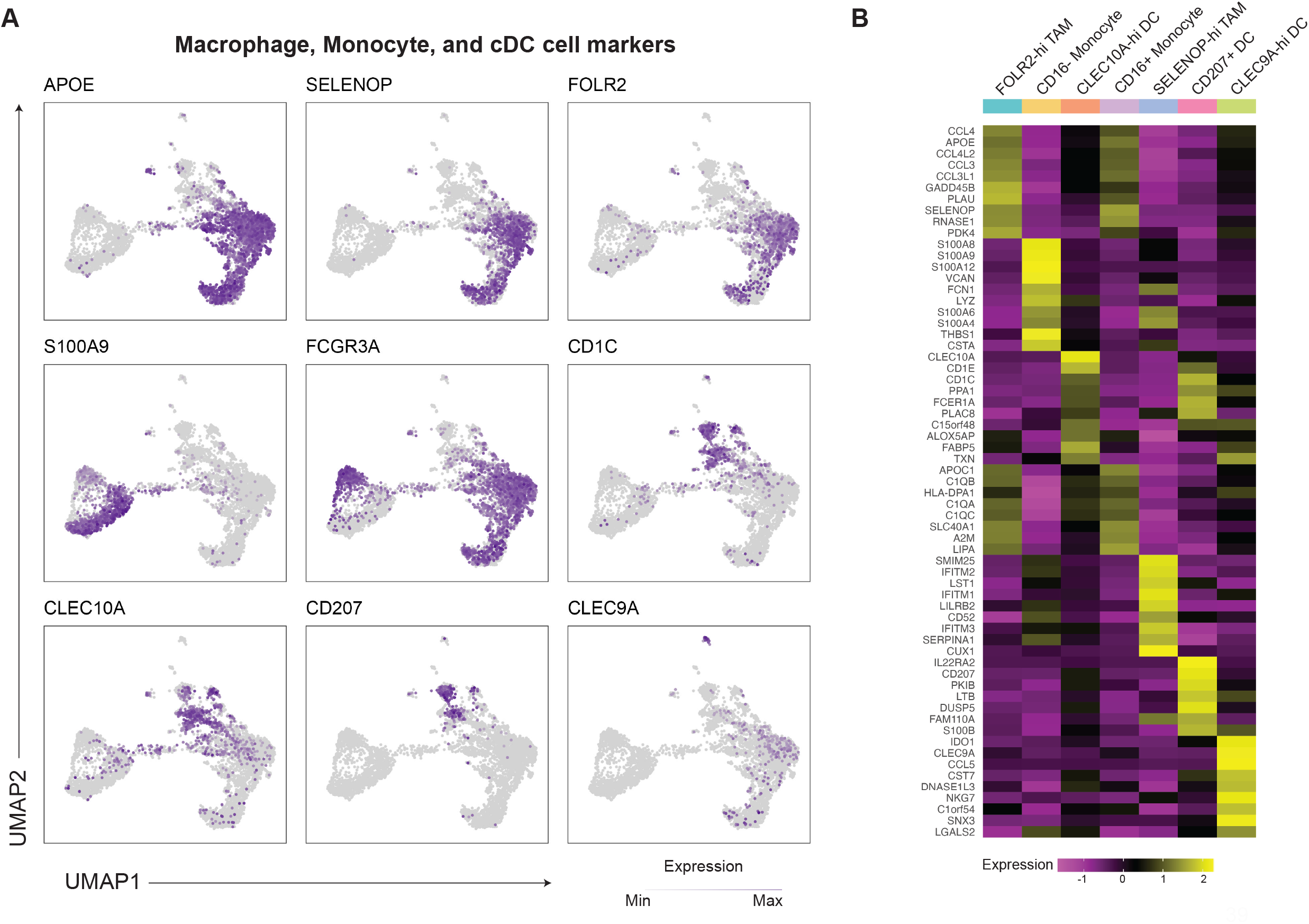
Related to Figure 4. A) Expression UMAP plots of macrophage, monocyte, and dendritic cells from all tumors, representing expression of marker genes used to identify myeloid subtypes. B) Heatmap of scaled and normalized average expression of top 10 differentially expressed genes between myeloid cell clusters.

